# Improving variant calling using population data and deep learning

**DOI:** 10.1101/2021.01.06.425550

**Authors:** Nae-Chyun Chen, Alexey Kolesnikov, Sidharth Goel, Taedong Yun, Pi-Chuan Chang, Andrew Carroll

## Abstract

Large-scale population variant data is often used to filter and aid interpretation of variant calls in a single sample. These approaches do not incorporate population information directly into the process of variant calling, and are often limited to filtering which trades recall for precision. In this study, we develop population-aware DeepVariant models with a new channel encoding allele frequencies from the 1000 Genomes Project. This model reduces variant calling errors, improving both precision and recall in single samples, and reduces rare homozygous and pathogenic clinvar calls cohort-wide. We assess the use of population-specific or diverse reference panels, finding the greatest accuracy with diverse panels, suggesting that large, diverse panels are preferable to individual populations, even when the population matches sample ancestry. Finally, we show that this benefit generalizes to samples with different ancestry from the training data even when the ancestry is also excluded from the reference panel.

## 1 Background

Variant calling [1–4] identifies the positions in an individual genome which differ from a reference or population, and is used to characterize a single sample or build large research cohorts [5, 6]. Variant calling is non-trivial, because of sequencing errors, systematic errors in mapping to repetitive and variable regions [7], and imbalanced sampling of alleles needed to identify a heterozygous variant from a homozygous one.

Variant calling can be improved by jointly genotyping multiple samples together [8–10], but the raw sequence data for a cohort is not always available, and this process is computationally expensive. Instead, large-scale reference panels from a wide range of populations can provide similar information [5, 6]. Recent studies use such information to improve alignment accuracy and reduce biases in alignment [11–13], but there has been little work to incorporate population data with variant calling.

Because far more variants are transmitted than arise de novo, real variants in a population tend to recur at various frequencies [14], while false positives are often either not seen elsewhere in a population, or are seen with a consistent signature [15]. Researchers use this knowledge to filter variant calls, often with rules which lose recall for a gain in precision [16]. More sophisticated machine-learning methods to filter are used in larger cohorts, such as gnomAD, but these also trade recall for precision and also only operate on variant calls and summary information [5].

We reason that including population-level information at an earlier stage in variant calling, when the full read-level data is available, might allow for more effective use of population data. To do this, we adapted DeepVariant [2], which represents BAM information as a multi-dimensional pileup and uses a Convolutional Neural Network (CNN) to call variants. Because DeepVariant learns the features important for variant classification directly from the data, it allows us to feed in the population allele information as an additional channel (Figure 1).

**Figure 1:**
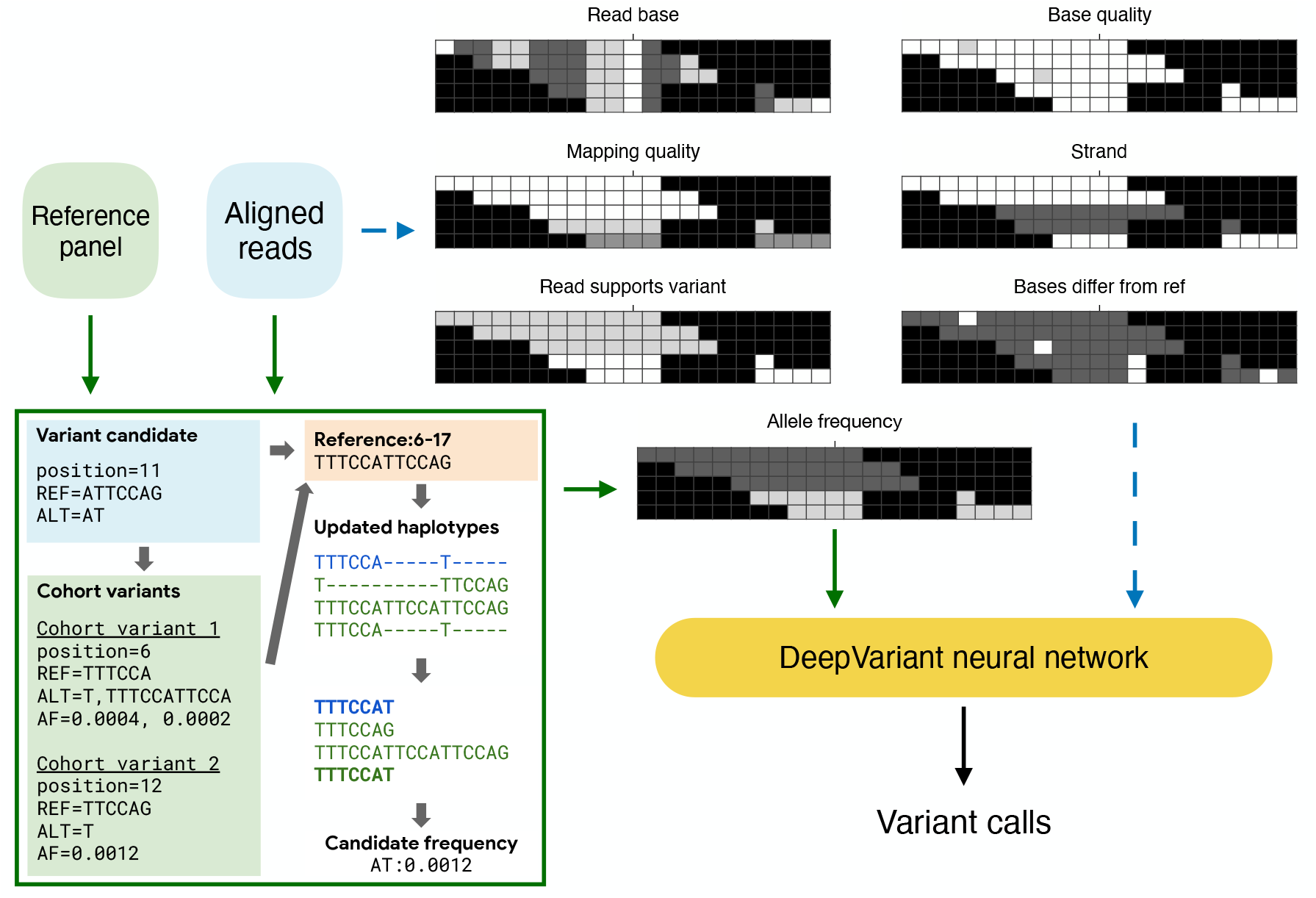
The population-aware DeepVariant (DeepVariant-AF) model. Dashed blue lines represent the typical population-agnostic DeepVariant approach, and the green lines show the data flow of the population-aware method. The green box shows the allele-matching algorithm to match variant alleles with a reference panel. This algorithm first queries cohort variants overlapped with the variant candidate and determines the window where haplotypes are updated. It then compares the haplotypes and updates the allele frequency of the matched ones

We trained population-aware models and compared them with the default DeepVariant v1.1 models which are agnostic of population information. The population-aware approach reduces the number of errors for all tested datasets, including WGS and WES reads, when using the allele frequencies from 1000Genomes. It also shows stronger error reduction efficacy for lower-coverage read sets. While traditional filtering approaches will increase precision at the expense of recall, we observe improvements to both precision and recall with this method.

When incorporating population data, it is also important for fairness and equity to understand how it changes the accuracy of methods for individuals with ancestries outside of those used in the development of the population resources. It is known that many genomic databases have collected more data for the European population than others [17–19]. We demonstrate that even using frequencies from a genetically distinct population, the population-aware model still performs similarly as the baseline. We find that a reference panel consisting of all ancestries in the 1000 Genomes Project (1000Genomes) outperforms a reference panel with only one of the 1000Genomes population groups, even when that population matches the sample being called. This implies that maximizing the diversity of ancestries in population resources has the potential to improve variant calling for all populations.

The Genome in a Bottle (GIAB) truth sets used to train DeepVariant are from European, Ashkenazi, and Asian ancestry [20]. To assess whether the addition of the reference panel information improves variant calling for populations outside of the populations represented in training, we use high quality PacBio HiFi data from the Human Genome Structural Variation Consortium for an individual of Puerto Rican ancestry as an evaluation set [21]. We show that an Illumina model using the reference panel has superior concordance with the highly accurate PacBio HiFi variant calls compared to an Illumina model without the reference panel.

## 2 Results

### 2.1 Population information improves variant calling performance

DeepVariant converts input from a BAM file into a pileup image with 6 channels, representing 1) bases, 2) base qualities, 3) mapping quality, 4) strand, 5) supports variant, and 6) base differs from reference. We modified DeepVariant v1.1 to take an additional input channel, the allele-frequency (AF) of the variant [22] (Figure 1). We trained DeepVariant models with and without the AF channel with the testing samples held out.

We assessed the variant calling results from the population-aware DeepVariant model (DeepVariant-AF), DeepVariant, GATK [4], Octopus [23] and Strelka2 [24]. We first compared the whole-genome sequencing (WGS) variant calling accuracy for sample HG003, sequenced with 35x coverage from the PrecisionFDA v2 Truth Challenge [25], using the latest GIAB v4.2.1 truth set [26] (Figure 2a and Table S1). HG003 is not used in the training of these DeepVariant models, and so acts as an independent holdout to evaluate their quality.

**Figure 2:**
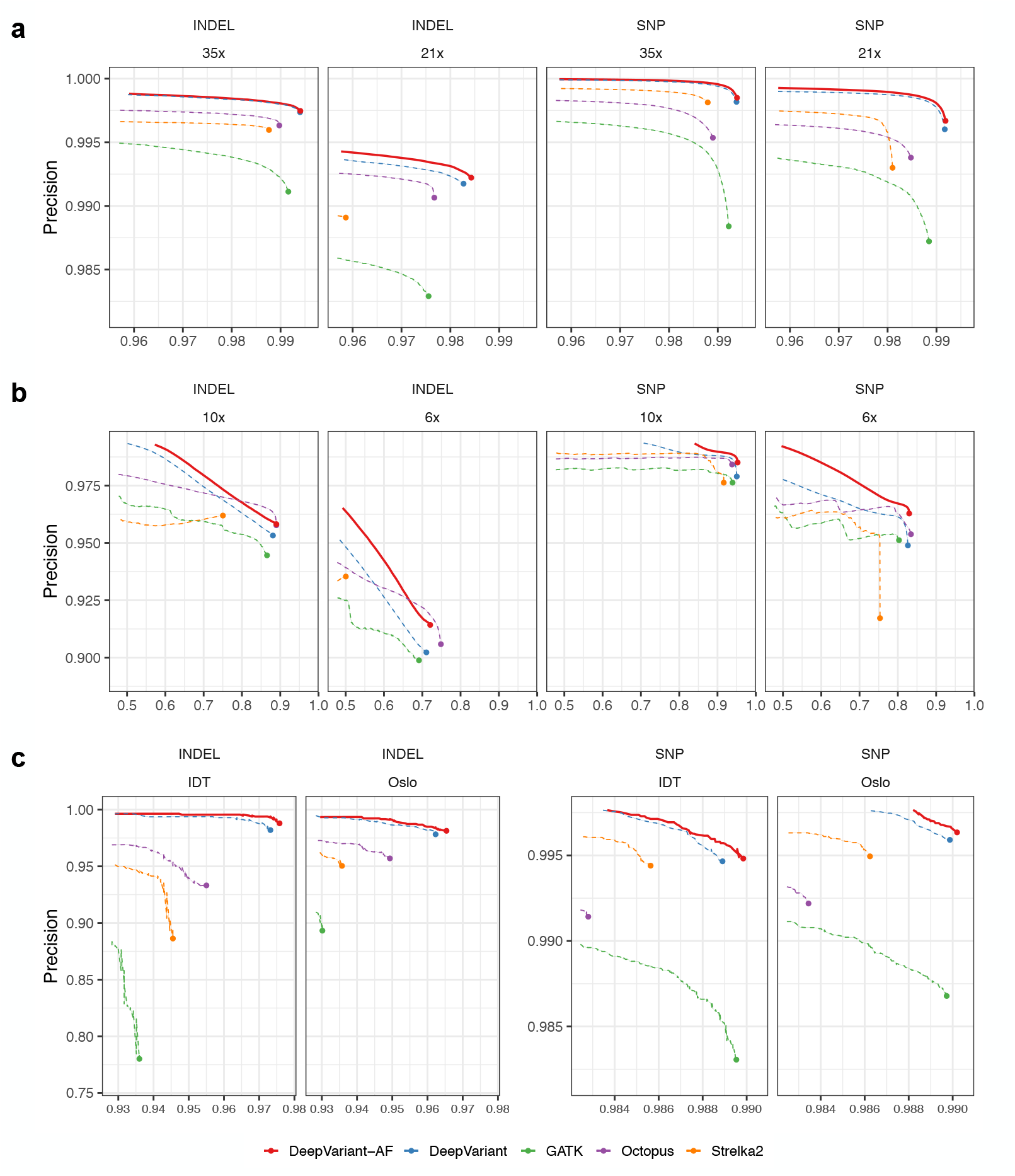
Variant calling accuracy using DeepVariant-AF and other methods. All datasets are from HG003. (a) High-coverage WGS datasets, (b) Low-coverage WGS datasets, (c) WES datasets. WGS results are evaluated using the GIAB v4.2.1 truth set (GRCh38) in the high-confidence regions. WES results are evaluated using the GIAB v 4.2.1 truth set (GRCh37) in the high-confidence regions that are captured

DeepVariant-AF has superior accuracy than all other methods in precision, recall and F1 score for both SNPs and indels. It has an overall error reduction of 1,499 (4.8%) compared to the second-best method (DeepVariant). Notably, DeepVariant-AF improves SNP precision from 0.9982 to 0.9985, equivalent to an error reduction of 1,068 (17.7%) variants. When compared to GATK, Octopus and Strelka2, DeepVariant-AF has error reductions of 44,009 (59.9%), 29,543 (50.0%) and 25,145 (46.0%) variants (SNPs and indels combined) respectively.

We then down-sampled the HG003 reads from 35x to 21x to evaluate the performance of the variant callers with lower-coverage datasets (Figure 2a). DeepVariant-AF demonstrates a larger improvement in accuracy over other methods. For example, DeepVariant-AF has an error reduction of 3,788 (7.0%) variants over the second-best method (DeepVariant). Similar to using the 35x read set, DeepVariant-AF shows the strongest improvement to reduce false-positive SNPs, improving precision from 0.9960 to 0.9967, equivalent to 2,202 (16.7%) errors. When further down-sampling the reads to 10x and 6x, DeepVariant-AF remains to be the method with the highest overall accuracy (Figure 2b). We then ran the Minimac4 imputation method [27] for all population-agnostic results and showed that DeepVariant-AF outperformed all imputation-based approaches (Note S1 and Table S1). We hypothesize that DeepVariant-AF is able to leverage the population information better when the sequence-based evidence gets weaker at lower coverages.

We further evaluated the performance of the models using two whole-exome sequencing (WES) datasets from a recently released set of genome and exome data for HG003 [28] (Figure 2c, Tables S2 and S3). Both datasets were aligned to GRCh37 and evaluated using the GIAB v4.2.1 truth set. For both WES datasets, DeepVariant-AF has the fewest overall errors among all tested callers. Compared to the second-best method (DeepVariant), it has overall error reduction levels of 8.1% (38 out of 469) for the IDT dataset and 6.4% (31 out of 487) for the Oslo dataset. Compared to other callers, DeepVariant-AF reduces 35.2% to 60.4% of the errors.

### 2.2 How does population information affect the model?

Intuitively, population information helps DeepVariant decide whether to make a call based on the commonness of a variant, especially for cases where the variant calling confidence levels are low. With a population-aware model, a variant caller should be more likely to make a positive variant call for a candidate with high allele frequency, and is less likely to make a call when seeing a rare candidate variant.

To understand the influence of allele frequencies in the model, we assessed the accuracy of DeepVariant-AF and other variant callers for common (allele frequency >0.01) and rare (allele frequency ≤0.01) variants using the 35x HG003 WGS dataset (Figure 3a and Section 4.5). The DeepVariant-AF shows substantial improvement over GATK and Strelka2, reducing 43.3-56.9% errors for common variants and 36.5-83.9% errors for rare variants. DeepVariant-AF also outperforms DeepVariant for both common and rare variants, reducing 892 (4.7%) and 931 (13.7%) errors respectively. There is enriched error reduction for false-negative common variants and false-positive rare variants by including population information in DeepVariant (Table S4 and S5).

**Figure 3:**
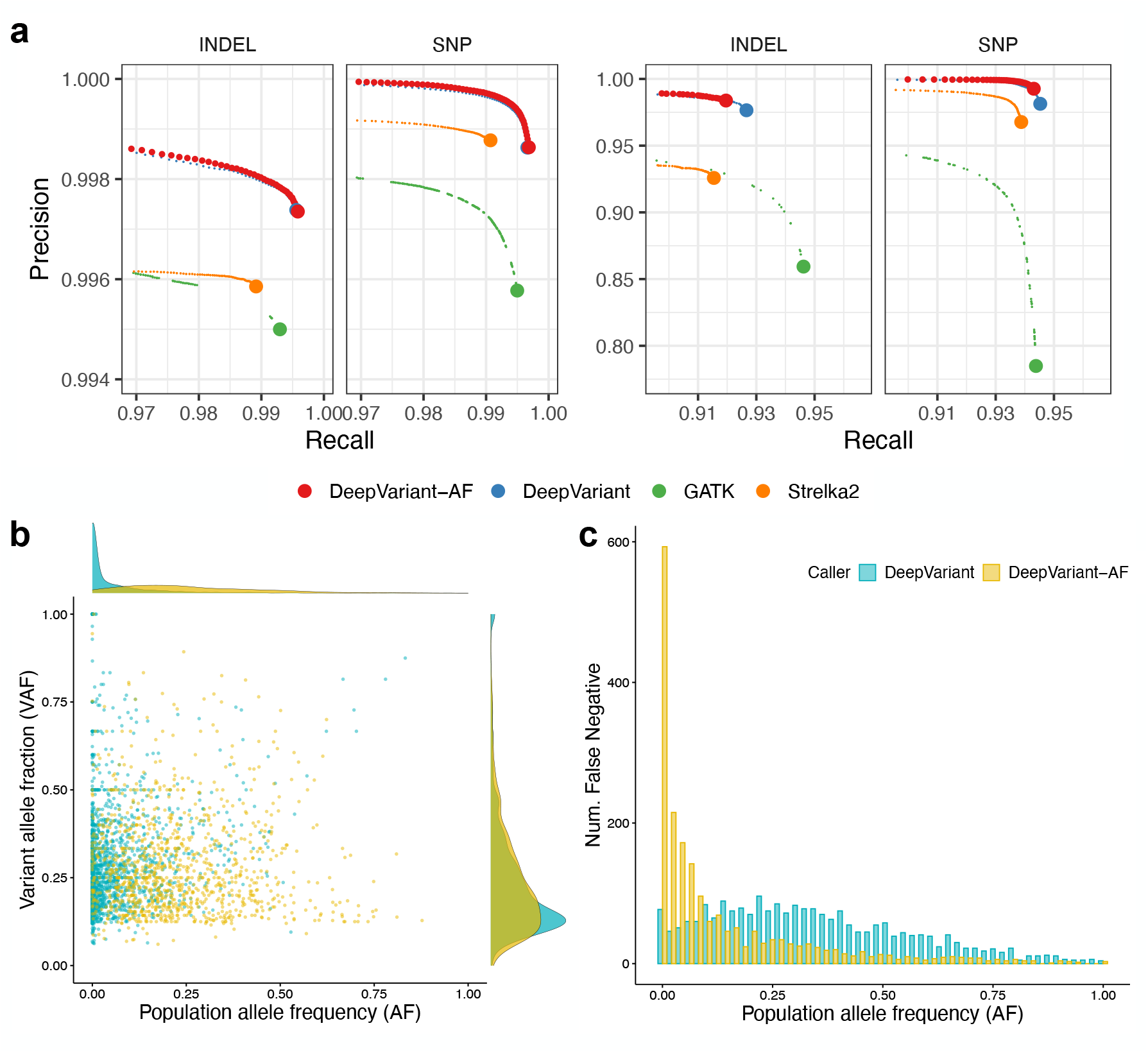
HG003 WGS variant calling results, annotated with 1000Genomes allele frequencies. (a) Variants are stratified by commonness. Left: common variants (allele frequency >0.01), right: rare variants (allele frequency ≤0.01). (b,c) Caller-specific errors by DeepVariant and DeepVariant-AF using 35x HG003 WGS data. Errors specific to Deep-Variant are considered to be population-resolved, and the others are considered to be population-induced. (b) false positives (DeepVariant: 2,085, DeepVariant-AF: 1,070), (c) false negatives (DeepVariant: 2,284, DeepVariant-AF: 1,952)

We also measured the recall for variants that appeared in the GIAB v4.2.1 truth set but had zero allele frequencies in 1000Genomes. Compared to the default DeepVariant model, DeepVariant-AF has a slightly lower recall but the difference was marginal (Figure S1). The recall of zero-frequency variants using all variant callers (71.4%-83.7% for SNPs and 88.1%-89.8% for indels) is substantially lower than the recall of all variants, but it can be strongly improved using PacBio Hifi reads ( Note S2). This implies many of the zero-frequency variants are hard to genotype using Illumina reads, and may not be novel mutations relative to samples in reference panels. In the future, reference panels utilizing high-quality long reads [29–31] will likely provide better allele frequency estimates and improve the population-aware model performance.

We further designed an analysis framework to assess errors specific to each variant calling method (Section 4.6). We compared the DeepVariant and DeepVariant-AF methods and identified false-positive and false-negative variants specific to each method. Variants specific to DeepVariant were “rescued” by population information and thus considered as “population-resolved”; whereas variants specific to DeepVariant-AF were considered to be induced by the population-aware model, likely due to the network adjustments when training using allele frequency data. We excluded errors common to both methods, since they were viewed as ones more difficult to resolve without major changes in the pipeline, such as the upstream data processing and sequencing methods.

We first examined the relationship between population allele frequency (AF) and variant allele fraction (VAF), which is the fraction of reads supporting an alternate allele in a given sample, of each false-positive call. There is an observable distinction between the population-induced group and the population-resolved group in the VAF-AF plots (Figure 3b). Among the population-resolved false-positive errors, more than one half (54.0%, or 1,125 out of 2,085) are rare among the 1000Genomes samples, whereas there are only 2.5% (49 out of 1,952) rare variants among the population-induced false positives.

We then investigated false-negative errors, as shown in Figure 3c. Variant allele fraction for false negatives are not always available because many false negatives are not identified as a variant candidate due to reasons including low read coverage, incorrect mapping or insufficient sensitivity in variant candidate discovery. Thus, we only evaluated the allele frequency distribution for false negatives. The number of erroneous common variants differs notably between the methods. Among all population-resolved false negatives, 96.6% (2,207 out of 2,284) are common variants. In contrast, only 30.1% (588 out of 1,952) of the population-induced false negatives are common. With the population knowledge provided in the AF channel, DeepVariant adjusts its variant calls according to the commonness of a variant and makes improvements in both precision and recall.

### 2.3 Assessing biases using different 1000Genomes populations

It is important to understand if the inclusion of population information reduces Deep-Variant’s performance for populations that are not well represented, especially when they have a large genomic difference with the reference panel. We first note that Ashkenazi Jewish, the ethnicity of the HG003, is not among the 26 ethnicities collected by 1000Genomes. Using a testing sample not in the reference panel reduces the risk of bias. Second, we ran inference on the population-aware model using reference panels of allele frequencies. We split the 1000Genomes sample into five groups based on the superpopulation labels (African, AFR; Admixed American, AMR; East Asian, EAS; European, EUR; South Asian, SAS) and calculated allele frequencies for each super-population.

We evaluated the accuracy using the 35x WGS HG003 dataset (Table 1). As described above, using the frequencies from the entire 1000Genomes demonstrates superior accuracy compared to the population-agnostic DeepVariant model. When inferencing using ancestry-specific frequencies, all DeepVariant-AF models outperform the baseline for SNPs, but underperform for indels. When considering the overall number of errors, only the model inferred with EAS frequencies calls more errors than the baseline, but the deficit (494, or 1.6% of the baseline) is small.

**Table 1:**
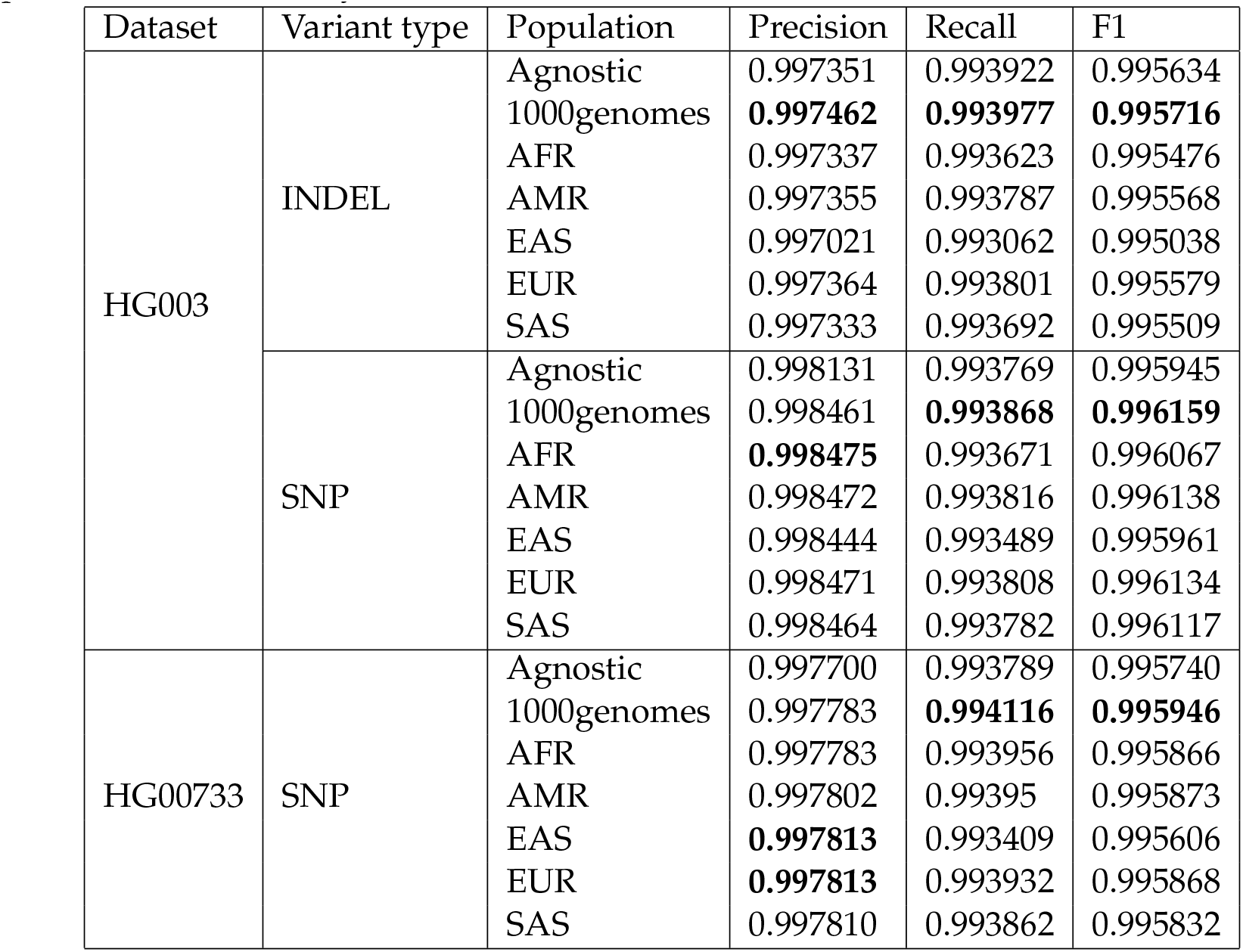
Variant calling accuracy when inferring using 35x WGS data from HG003 and 30x WGS data from HG00733. Methods: default DeepVariant (*Agnostic*), population-aware DeepVariant using allele frequencies from the entire 1000Genomes (*1000genomes*) and five 1000Genomes superpopulations (*AFR*, *AMR*, *EAS*, *EUR* and *SAS*). Higher values correspond to better accuracy

| Dataset | Variant type | Population | Precision | Recall | F1 |
| --- | --- | --- | --- | --- | --- |
| HG003 | INDEL | Agnostic | 0.997351 | 0.993922 | 0.995634 |
|  |  | 1000genomes | **0.997462** | **0.993977** | **0.995716** |
|  |  | AFR | 0.997337 | 0.993623 | 0.995476 |
|  |  | AMR | 0.997355 | 0.993787 | 0.995568 |
|  |  | EAS | 0.997021 | 0.993062 | 0.995038 |
|  |  | EUR | 0.997364 | 0.993801 | 0.995579 |
|  |  | SAS | 0.997333 | 0.993692 | 0.995509 |
|  | SNP | Agnostic | 0.998131 | 0.993769 | 0.995945 |
|  |  | 1000genomes | 0.998461 | **0.993868** | **0.996159** |
|  |  | AFR | **0.998475** | 0.993671 | 0.996067 |
|  |  | AMR | 0.998472 | 0.993816 | 0.996138 |
|  |  | EAS | 0.998444 | 0.993489 | 0.995961 |
|  |  | EUR | 0.998471 | 0.993808 | 0.996134 |
|  |  | SAS | 0.998464 | 0.993782 | 0.996117 |
| HG00733 | SNP | Agnostic | 0.997700 | 0.993789 | 0.995740 |
|  |  | 1000genomes | 0.997783 | **0.994116** | **0.995946** |
|  |  | AFR | 0.997783 | 0.993956 | 0.995866 |
|  |  | AMR | 0.997802 | 0.99395 | 0.995873 |
|  |  | EAS | **0.997813** | 0.993409 | 0.995606 |
|  |  | EUR | **0.997813** | 0.993932 | 0.995868 |
|  |  | SAS | 0.997810 | 0.993862 | 0.995832 |

We also compared the performance of using different superpopulation allele frequencies and observed that using frequencies from a genetically closer population usually resulted in higher variant calling accuracy. Using EUR frequencies reduces 1,700 (5.3%) more variants than using EAS frequencies, echoing the estimation that Ashkenazi Jewish is genetically closer to the European populations and is farther from East Asian and African populations [5, 32–34]. We point out that using 1000Genomes frequencies from all populations results in the lowest number of errors among all population-aware results, suggesting an advantage to using a diverse population than finding a genetically similar group. This finding echoes our previous statement that we anticipate the population-aware variant calling model to improve further with larger-scaled and more diverse population callsets.

### 2.4 Silver-standard truth set for HG00733

Genome-in-a-bottle (GIAB) truth variant sets provide gold standards to benchmark variant callers, but until now there are only three samples (HG002-HG003-HG004, the Ashkenazi trio) with curated calls in difficult-to-map regions added in the v4.2.1 release [26]. Further, the samples are from the same ancestry, making it challenging to perform a generalized benchmarking considering the genetic diversity of the human population. To deal with this difficulty, it is desirable to have other high-quality variant sets from non-GIAB samples, preferably from ancestries not covered by GIAB. Thus, we called variants using the DeepVariant PacBio model with 32x high-coverage PacBio HiFi reads [35] for HG00733, a Puerto Rican (labelled as PUR under the AMR superpopulation in 1000Genomes) sample. The DeepVariant PacBio model has a SNP F1 score higher than 99.9% and is one of the most accurate models using PacBio HiFi data [26]. We used the DeepVariant HG00733 PacBio SNP calls as a “silver-standard” truth set and benchmarked the performance for models using Illumina reads. We used 30x Illumina WGS reads sequenced by the New York Genome Center [36] to test all HG00733 models. Because the 1000Genomes has a collection of PUR samples, we excluded all PUR samples and recalculated allele frequencies for both 1000Genomes and the AMR superpopulation.

DeepVariant-AF has a higher SNP F1 (0.9950) than DeepVariant (0.9948) and other variant callers (GATK: 0.9912, Octopus: 0.9918, Strelka2: 0.9923) (Figure 4), reducing 1,527 to 26,045 variants (4.4% to 43.8%). Similar to the finding using HG003, DeepVariant-AF performs strongly with a down-sampled (18x) read set by reducing 4,423 to 44,537 (9.5% to 52.3%) erroneously called SNPs. The lead is observed for even lower coverage datasets (10x and 6x). Though the accuracy difference between DeepVariant-AF and Octopus is small at 6x, DeepVariant-AF still outperforms by an error reduction of 18,368 (3.5%) variants.

**Figure 4:**
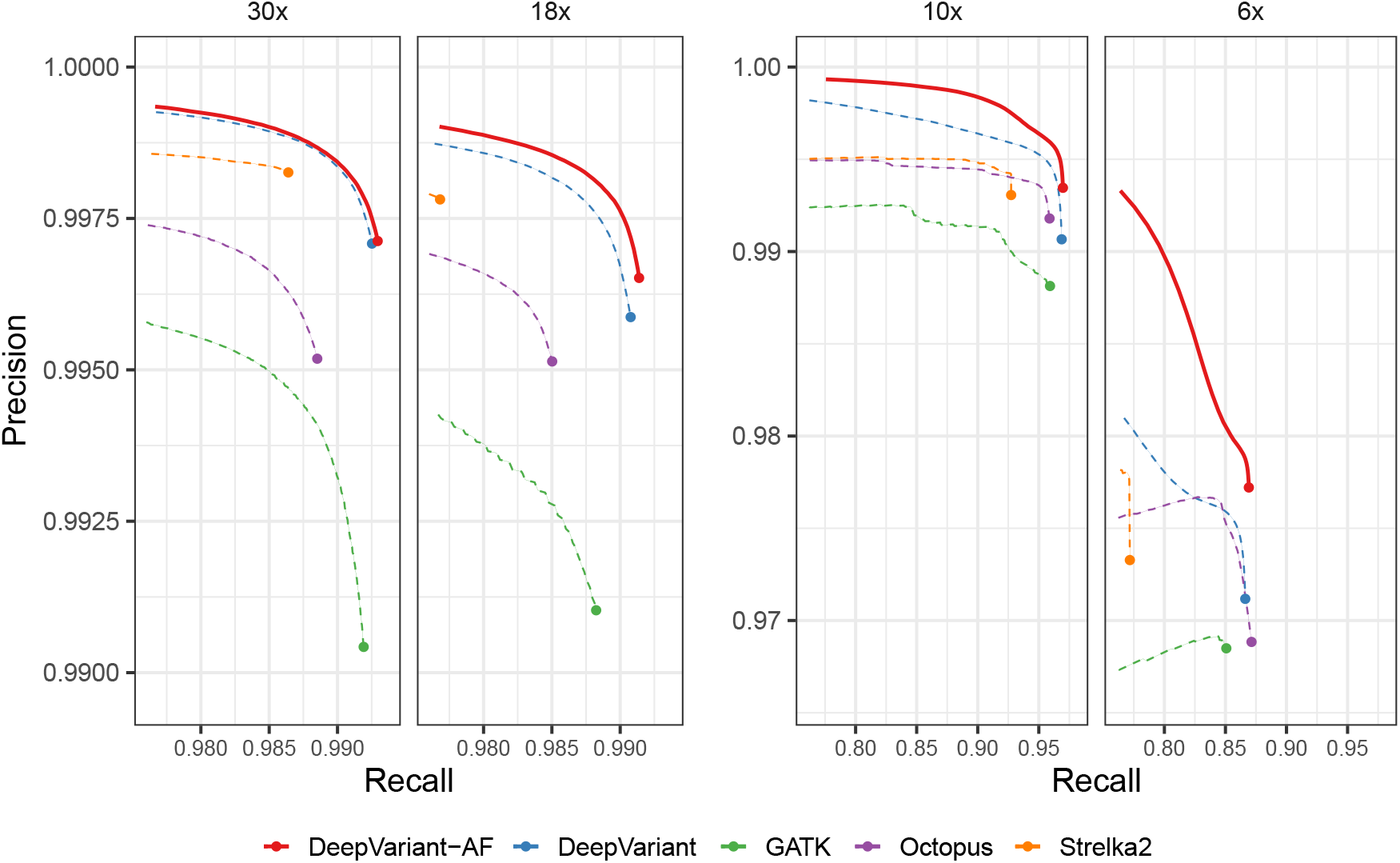
SNP calling accuracy using DeepVariant-AF and other methods using HG00733 WGS data. The results are compared to the PacBio-DeepVariant silver-standard truth set

We also tested the model using different superpopulation frequencies (Table 1). All but the EAS population-aware model has higher SNP F1 scores than the baseline. Using DeepVariant-AF and inferring using the EAS allele frequencies results in 878 (3.1%) more errors. All population-aware models, including EAS, outperform the baseline in precision and only EAS has a lower recall than the baseline (0.993409 vs. 0.993789). We note that all the tested DeepVariant-AF models outperform other non-DeepVariant methods in SNP F1 accuracy.

### 2.5 Population-aware models have a larger effect on the cohort level for rare variant calls

Variant calling is often applied to large scale cohorts to generate a population-level callset across many samples [37]. In large cohorts, rare variants present a unique opportunity to discover variant associations with large effect sizes, such as loss-of-function variants [38, 39]. These analyses aggregate the signal from several variants in the same gene or pathway [40]. However, this analysis must also contend with the impact of false positive calls.

Because the population-aware model has a higher precision for rare variants, and because rare false positive calls aggregate across many samples at the cohort level, we reasoned that the improved accuracy of the population-aware model could be larger for rare variants.

To test this, we generated cohort-wide calls of the recent whole genome sequencing of the 1000Genomes using both the DeepVariant v1.1 out-of-the-box WGS model, and the allele frequency-aware model. To investigate the effect on rare variants, we looked at variant metrics for calls present in only 1 sample (singleton), as well as those in a small number of samples.

We observed a large reduction in rare homozygous variants (Figure 5a), which can have a large effect on analysis of recessive loss-of-function variants. Similarly, we saw a reduction in the number of rare variants which are known to be pathogenic or likely pathogenic in Clinvar [41] (Figure 5b). The increased precision for rare variants in a single sample suggests that this reduction may be achieved by reducing the number of false positive calls, which is supported by an increase in the transition:transversion (Ti:Tv) ratio, an indirect measure of call quality, for homozygous rare variants (Figure 5c) and heterozygous rare variants (Figure 5d), with a more pronounced improvement for rare homozygous variants.

**Figure 5:**
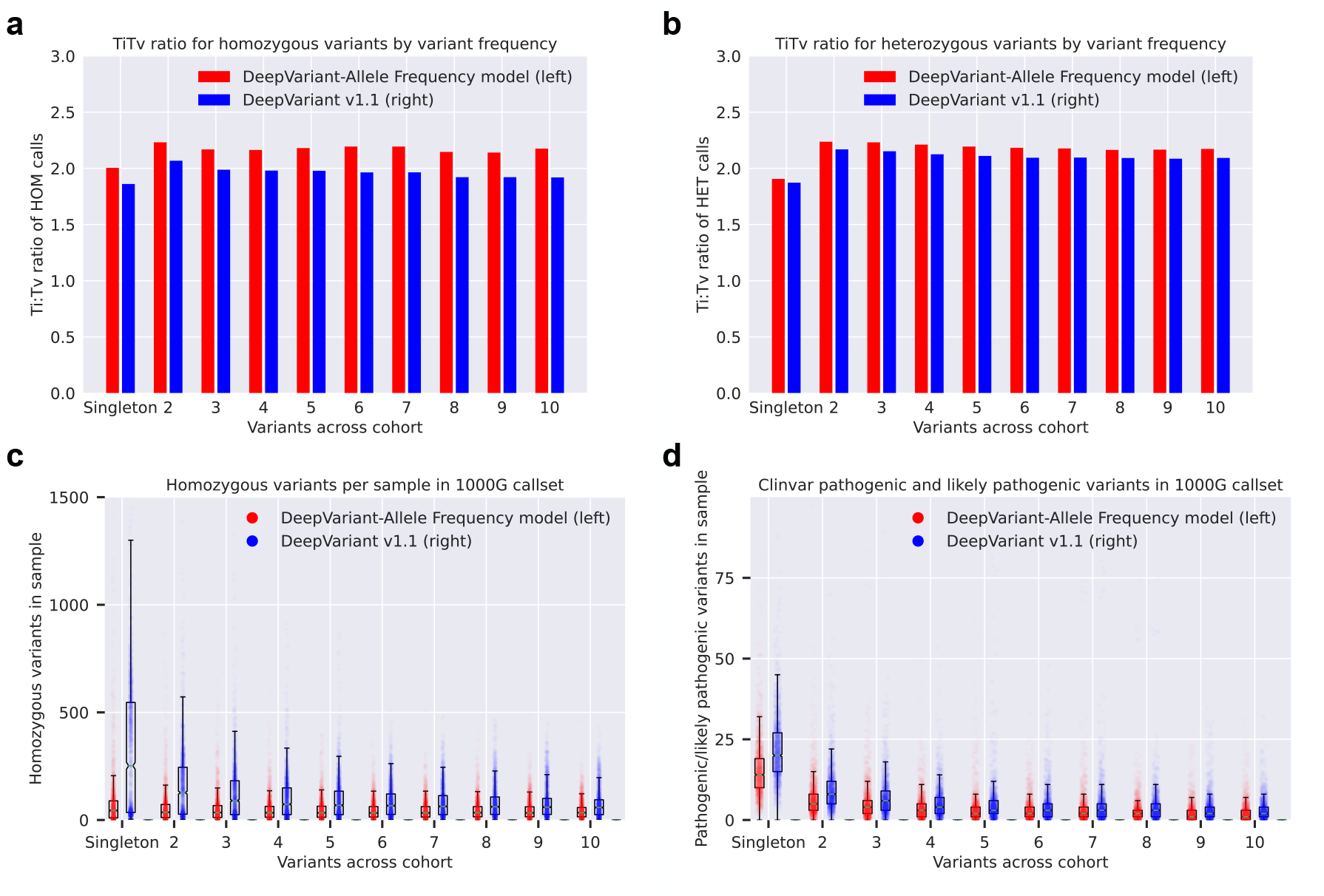
Cohort-level rare variant metrics in the 1000genomes using DeepVariant-AF and DeepVariant v1.1. Calls in each plot are stratified by the frequency of calls in a sample, ranging from a call present in only one sample (singleton) to a call in 10 different samples. (a) The number of homozygous variant calls per sample (each dot is one 1000g sample). (b) The number of Clinvar pathogenic or likely pathogenic variants per sample (each dot is one 1000g sample). (c) The Ti:Tv ratio for calls by frequency for homozygous variant calls, averaged across all samples. (d) The Ti:Tv ratio for calls by frequency for heterozygous variant calls, averaged across all samples

## 3 Discussion

We designed a new population-aware DeepVariant model which can incorporate both base- and read-level information with the population information. We find that population-aware models reduce error rates compared to other state-of-the-art variant calling methods. The relative advantage of the population-aware models increase at lower coverage, suggesting that population information is most valuable in difficult examples, where read-level information alone may not be sufficient for confident calling. In population sequencing projects, this finding could be relevant to the question of whether to sequence more individuals at lower coverage, or fewer at a high coverage. When sequencing for a species without a reference panel, it is possible that sequencing more, diverse individuals at lower coverage could still retain comparable accuracy to traditional methods which do not incorporate population information in calling.

We evaluate potential biases introduced by population information in variant calling by comparing population-aware models that use allele frequencies from different 1000Genomes superpopulation. This experiment simulates a scenario where the tested sample is genetically distinct from the reference panel. Only one population-aware method (inferred with EAS frequencies) underperforms the baseline in total number of errors, but with a small deficit. Furthermore, using allele frequencies calculated from the entire 1000Genomes outperforms population-specific methods. This finding implies that a diverse population can provide more benefits than using a homogeneous one, even when the homogeneous population is more genetically similar with the tested sample. This finding may inform efforts to build population or country-specific resources. Increasing the number of samples for a given population will improve accuracy for that population, but the inclusion of samples from diverse populations will also improve the resource. We believe that the accuracy of the population-aware model can further improve with a larger and more diverse population callset in the future, reinforcing the benefit of collaboration between nation-scale efforts.

We provide an additional “silver-standard” SNP set for a Purto Rican sample, HG00733, a population not present in the labeled training data. We used high-coverage PacBio HiFi reads and an accurate DeepVariant PacBio model to generate this high-quality call set. This method can provide high-confidence SNP calls for non-GIAB samples and increase population diversity when assessing variant calling results. Similar to the results using HG003 data, we show that the proposed model has strong performance compared to the baseline, and only suffers slight loss of accuracy when inferred using a distinct population. When more high-coverage PacBio HiFi data become available in the future [29–31], the high-quality calls generated by DeepVariant can provide a more diversified dataset for variant calling benchmarking and downstream analysis.

The largest differences that we observe with the population-aware models occur at the cohort level, with potentially larger implication for the analysis of rare variants within these cohorts. We see substantial reductions in the number of both rare homozygous variants and variants that are annotated as pathogenic or likely pathogenic in Clinvar. This may occur by reducing false positives, and by making heterozygous calls more likely when a rare variant could plausibly be heterozygous or homozygous. Increasing the precision for these rare variants across the cohort could increase the statistical signal of rare variant binning approaches, and improve the discovery of rare impact associations relevant to phenotypic traits.

We also notice that all tested Illumina models performed poorly on the zero-frequency variants, regardless of using population information or not. By analyzing the variants with PacBio reads, we point out many zero-frequency variants in 1000Genomes located in difficult-to-map regions, but likely not genetically novel in the population. This suggests that the power of population-aware methods should increase as large panels of long-read population data become available.

## 4 Methods

### 4.1 Model training

We trained the model following the procedure described in [2], with additional Illumina WGS datasets included [28]. Variants in chromosomes 1 to 19 are used as the training examples, and those in chromosome 21 and 22 are used for tuning. Variants in chromosome 20 are never used in the training process.

### 4.2 Datasets

The model is evaluated using the GIAB v4.2.1 truth set for HG003 across the whole genome [26]. We also generated another high-quality SNP set using DeepVariant v0.10 and HG00733 PacBio HiFi data [35] across the whole genome. We used the intersection of high-confidence regions of HG002, HG003, and HG004 (GIAB v4.2.1) as the high-confidence regions for the HG00733 SNP set. The read sets used for experiments are listed in Table 2 and the read sets for supporting experiments are provided in Table 3.

**Table 2:** Testing datasets

| Sample | Ethnicity | Truth variant | Dataset |
| --- | --- | --- | --- |
| HG003 | Ashkenazi Jewish | v4.2.1 (GRCh38) | 35x Illumina WGS [26] |
| HG003 | Ashkenazi Jewish | v4.2.1 (GRCh37) | 100x Illumina WES [28]<br>300x Illumina WES [20] |
| HG00733 | Puerto Rican | DeepVariant v0.10<br>PacBio SNP calls (GRCh38) | 30x Illumina WGS [36] |

**Table 3:** Other datasets used in this study

| Sample | Ethnicity | Dataset |
| --- | --- | --- |
| HG003 | Ashkenazi Jewish | 35x PacBio HiFi [26] |
| HG00733 | Puerto Rican | 32x PacBio HiFi [35] |

### 4.3 Allele matching algorithm

When incorporating population information in DeepVariant, we need to match a variant candidate with a cohort variant. However, this is not a straightforward task since a variant can be represented in multiple formats [3, 42, 43]. A common approach is to normalize variants, such as using `bcftools norm` [44], but that’s not sufficient for complicated cases.

We designed an algorithm that constructed local haplotypes and performed precise allele matching (Figure 1, inset). The algorithm starts with querying all cohort variants *V*_*C*_ overlapped with a window [*v*_*start*_, *v*_*end*_), where *v*_*start*_ and *v*_*end*_ are the starting and ending positions of a variant candidate *v* respectively. The queried cohort variants and the candidate variant form set *V* ≡ *v* ⋃ *V*_*C*_. Then the window is extended to the smallest starting position and the largest ending position within *V*, as [*V*_*start*_, *V*_*end*_), where *V*_*start*_ ≡ min(*u*_*start*_)∀*u* ϵ *V* and *V*_*end*_ ≡ *max*(*end*_*w*_) ∀*w* ϵ *V*. Local reference haplotype is queried from the reference genome in window [*V*_*start*_, *V*_*end*_). For each variant allele in *V*, we construct its local allele haplotype. If there’s a perfect match between a cohort allele haplotype and a candidate allele haplotype, the allele frequency of the cohort allele is added to an allele frequency dictionary, using the alternate allele of the candidate variant as its key. Afterwards, DeepVariant looks up the dictionary to update the allele frequency of each read that overlaps with the candidate variant.

### 4.4 Allele frequency channel for DeepVariant

To make full advantages of the CNN-based classifier of DeepVariant, allele frequencies need to be encoded in pileup images. We apply a logarithmic transformation to gain resolution for low-frequency signals. For each variant candidate, an additional *allele frequency channel* is added to the pileup image. In this channel, a read is colored by the transformed frequency of its allele at the variant candidate position. A read can carry multiple alternate alleles with different frequencies, so its color intensity may vary across pileup images, where the variant candidates differ. An alternative method to encode allele frequencies is to include the information as features in the fully-connected layers [45], but this approach sacrifices the capability to incorporate allele frequencies with base- and read-level information and thus is not adopted.

To enable the allele frequency channel, users need to enable flag --`use allele frequency` and provide DeepVariant cohort variants in VCF format with flag --`population vcfs<vcf>`.

### 4.5 Stratifying variants by commonness

To measure precision, we matched the called variants with the 1000Genomes reference panel and annotated allele frequencies using the allele matching algorithm. Similarly, we annotated the allele frequency of GIAB v4.2.1 truth variants and measured recall. We excluded multi-allelic variants where one allele is common and the other is rare. We didn’t perform this analysis for results from Octopus because the variants were represented differently.

### 4.6 Model-specific error analysis

We compared actual variant calls with GIAB v4.2.1 truth variants. Variants specific to actual calls are regarded as false positives, and those specific to the truth set are false negatives. We generated the false-positive and false-negative sets for two models, and obtained model-specific false positives and false negatives. For both sets, we applied the allele matching algorithm to annotate the allele frequency (AF) of the variants. For the false-positive sets, we extracted variant allele fractions (VAF) from the VCF files generated by DeepVariant.

### 4.7 1000Genomes frequencies from the DeepVariant-GLnexus pipeline

We used the 1000Genomes reference panel generated with the DeepVariant-GLnexus pipeline (v3) [9] for all population-aware experiments, including training and inferring the models. We filled the missing genotypes with the reference genotypes with `bcftools +missing2ref` to make sure all variants have the same denominator.

## 5 Availability of data and materials

### 5.1 Software

The DeepVariant source code is available at https://github.com/google/deepvariant under the BSD-3-Clause License. The pre-trained population-aware DeepVariant models are available at https://console.cloud.google.com/storage/browser/brain-genomics-publresearch/allele_frequency/pretrained_model_WGS (WGS) and https://console.cloud.google.com/storage/browser/brain-genomics-public/research/allele_frequency/pretrained_model_WES (WES).

### 5.2 Data

The 1000Genomes callset generated using the population-aware model is available at: https://console.cloud.google.com/storage/browser/brain-genomics-public/research/allele_frequency/1KGP/cohort_dv_af_glnexus. The PacBio-based HG00733 SNP set is available at https://console.cloud.google.com/storage/browser/brain-genomics-public/research/allele_frequency/HG00733_SNP_set. The VCF files used in this study are available at https://console.cloud.google.com/storage/browser/brain-genomics-public/research/cohort/1KGP/cohort_dv_glnexus_opt/v3_missing2ref (GRCh38) and https://console.cloud.google.com/storage/browser/brain-genomics-public/research/cohort/1KGP/cohort_dv_glnexus_opt/v3_GRCh37_missing2ref (GRCh37).

## 6 Ethics approval and consent to participate

Not applicable.

## 7 Consent for publication

Not applicable.

## 10 Acknowledgments

We thank Babak Alipanahi, Gunjan Baid, Daniel Cook, Alexander D’Amour, Hojae Lee, Cory McLean, Maria Nattestad and other colleagues at Google and Ben Langmead for their feedback on this manuscript and the project in general.

## 11 Authors’ contributions

NC, AK, PC and AC designed the method. NC, AK and PC implemented the software. NC, TY, PC and AC performed the experiment. NC, AK, SG, TY, PC and AC analyzed the results. NC, PC and AC wrote the manuscript. All authors read and approved the final manuscript.

## References

1. DePristo, M. A., Banks, E., Poplin, R., Garimella, K. V., Maguire, J. R., Hartl, C., Philippakis, A. A., Del Angel, G., Rivas, M. A., Hanna, M., et al. A framework for variation discovery and genotyping using next-generation DNA sequencing data. Nature genetics 43, 491 (2011).

2. Poplin, R., Chang, P.-C., Alexander, D., Schwartz, S., Colthurst, T., Ku, A., Newburger, D., Dijamco, J., Nguyen, N., Afshar, P. T., et al. A universal SNP and small-indel variant caller using deep neural networks. Nature biotechnology 36, 983–987 (2018).

3. Krusche, P., Trigg, L., Boutros, P. C., Mason, C. E., Francisco, M., Moore, B. L., Gonzalez-Porta, M., Eberle, M. A., Tezak, Z., Lababidi, S., et al. Best practices for benchmarking germline small-variant calls in human genomes. Nature biotechnology 37, 555–560 (2019).

4. Van der Auwera, G. A. & O’Connor, B. D. Genomics in the Cloud: Using Docker, GATK, and WDL in Terra (O’Reilly Media, 2020).

5. Karczewski, K. J., Francioli, L. C., Tiao, G., Cummings, B. B., Alföldi, J., Wang, Q., Collins, R. L., Laricchia, K. M., Ganna, A., Birnbaum, D. P., et al. The mutational constraint spectrum quantified from variation in 141,456 humans. Nature 581, 434–443 (2020).

6. 1000 Genomes Project Consortium et al. A global reference for human genetic variation. Nature 526, 68–74 (2015).

7. Li, H. Toward better understanding of artifacts in variant calling from high-coverage samples. Bioinformatics 30, 2843–2851 (2014).

8. Lin, M. F., Rodeh, O., Penn, J., Bai, X., Reid, J. G., Krasheninina, O. & Salerno, W. J. GLnexus: joint variant calling for large cohort sequencing. BioRxiv, 343970 (2018).

9. Yun, T., Li, H., Chang, P.-C., Lin, M. F., Carroll, A. & McLean, C. Y. Accurate, scalable cohort variant calls using DeepVariant and GLnexus. Bioinformatics 36, 5582–5589 (2020).

10. Poplin, R., Ruano-Rubio, V., DePristo, M. A., Fennell, T. J., Carneiro, M. O., Van der Auwera, G. A., Kling, D. E., Gauthier, L. D., Levy-Moonshine, A., Roazen, D., et al. Scaling accurate genetic variant discovery to tens of thousands of samples. BioRxiv, 201178 (2017).

11. Chen, N.-C., Solomon, B., Mun, T., Iyer, S. & Langmead, B. Reference flow: reducing reference bias using multiple population genomes. Genome biology 22, 1–17 (2021).

12. Rautiainen, M. & Marschall, T. GraphAligner: rapid and versatile sequence-to-graph alignment. Genome biology 21, 1–28 (2020).

13. Garrison, E., Sirén, J., Novak, A. M., Hickey, G., Eizenga, J. M., Dawson, E. T., Jones, W., Garg, S., Markello, C., Lin, M. F., et al. Variation graph toolkit improves read mapping by representing genetic variation in the reference. Nature biotechnology 36, 875–879 (2018).

14. Witherspoon, D. J., Wooding, S., Rogers, A. R., Marchani, E. E., Watkins, W. S., Batzer, M. A. & Jorde, L. B. Genetic similarities within and between human populations. Genetics 176, 351–359 (2007).

15. Abramovs, N., Brass, A. & Tassabehji, M. Hardy-Weinberg Equilibrium in the Large Scale Genomic Sequencing Era. Frontiers in Genetics 11, 210 (2020).

16. Pedersen, B. S., Brown, J. M., Dashnow, H., Wallace, A. D., Velinder, M., Tvrdik, T., Mao, R., Best, H. D., Bayrak-Toydemir, P. & Quinlan, A. R. Effective variant filtering and expected candidate variant yield in studies of rare human disease. BioRxiv (2020).

17. Sirugo, G., Williams, S. M. & Tishkoff, S. A. The missing diversity in human genetic studies. Cell 177, 26–31 (2019).

18. Martin, A. R., Kanai, M., Kamatani, Y., Okada, Y., Neale, B. M. & Daly, M. J. Clinical use of current polygenic risk scores may exacerbate health disparities. Nature genetics 51, 584–591 (2019).

19. McGuire, A. L., Gabriel, S., Tishkoff, S. A., Wonkam, A., Chakravarti, A., Furlong, E. E., Treutlein, B., Meissner, A., Chang, H. Y., López-Bigas, N., et al. The road ahead in genetics and genomics. Nature Reviews Genetics 21, 581–596 (2020).

20. Zook, J. M., Catoe, D., McDaniel, J., Vang, L., Spies, N., Sidow, A., Weng, Z., Liu, Y., Mason, C. E., Alexander, N., et al. Extensive sequencing of seven human genomes to characterize benchmark reference materials. Scientific data 3, 1–26 (2016).

21. Wenger, A. M., Peluso, P., Rowell, W. J., Chang, P.-C., Hall, R. J., Concepcion, G. T., Ebler, J., Fungtammasan, A., Kolesnikov, A., Olson, N. D., et al. Accurate circular consensus long-read sequencing improves variant detection and assembly of a human genome. Nature biotechnology 37, 1155–1162 (2019).

22. Carroll, A. & Chang, P.-C. Improving the Accuracy of Genomic Analysis with DeepVariant 1.0 https://ai.googleblog.com/2020/09/improving-accuracy-of-genomic-analysis.html. 2020. (accessed: 2020-12-11).

23. Cooke, D. P., Wedge, D. C. & Lunter, G. A unified haplotype-based method for accurate and comprehensive variant calling. Nature biotechnology, 1–8 (2021).

24. Kim, S., Scheffler, K., Halpern, A. L., Bekritsky, M. A., Noh, E., Källberg, M., Chen, X., Kim, Y., Beyter, D., Krusche, P., et al. Strelka2: fast and accurate calling of germline and somatic variants. Nature methods 15, 591–594 (2018).

25. Olson, N. D., Wagner, J., McDaniel, J., Stephens, S. H., Westreich, S. T., Prasanna, A. G., Johanson, E., Boja, E., Maier, E. J., Serang, O., et al. precisionFDA Truth Challenge V2: Calling variants from short-and long-reads in difficult-to-map regions. bioRxiv (2020).

26. Wagner, J., Olson, N. D., Harris, L., Khan, Z., Farek, J., Mahmoud, M., Stankovic, A., Kovacevic, V., Wenger, A. M., Rowell, W. J., et al. Benchmarking challenging small variants with linked and long reads. BioRxiv (2020).

27. Das, S., Forer, L., Schönherr, S., Sidore, C., Locke, A. E., Kwong, A., Vrieze, S. I., Chew, E. Y., Levy, S., McGue, M., et al. Next-generation genotype imputation service and methods. Nature genetics 48, 1284–1287 (2016).

28. Baid, G., Nattestad, M., Kolesnikov, A., Goel, S., Yang, H., Chang, P.-C. & Carroll, A. An extensive sequence dataset of gold-standard samples for benchmarking and development. bioRxiv (2020).

29. Ebert, P., Audano, P. A., Zhu, Q., Rodriguez-Martin, B., Porubsky, D., Bonder, M. J., Sulovari, A., Ebler, J., Zhou, W., Mari, R. S., et al. Haplotype-resolved diverse human genomes and integrated analysis of structural variation. Science 372 (2021).

30. Beyter, D., Ingimundardottir, H., Oddsson, A., Eggertsson, H. P., Bjornsson, E., Jonsson, H., Atlason, B. A., Kristmundsdottir, S., Mehringer, S., Hardarson, M. T., et al. Long-read sequencing of 3,622 Icelanders provides insight into the role of structural variants in human diseases and other traits. Nature Genetics 53, 779–786 (2021).

31. De Coster, W., Weissensteiner, M. H. & Sedlazeck, F. J. Towards population-scale long-read sequencing. Nature Reviews Genetics, 1–16 (2021).

32. Einhorn, Y., Weissglas-Volkov, D., Carmi, S., Ostrer, H., Friedman, E. & Shomron, N. Differential analysis of mutations in the Jewish population and their implications for diseases. Genetics research 99 (2017).

33. Xue, J., Lencz, T., Darvasi, A., Pe’er, I. & Carmi, S. The time and place of European admixture in Ashkenazi Jewish history. PLoS genetics 13, e1006644 (2017).

34. Carmi, S., Hui, K. Y., Kochav, E., Liu, X., Xue, J., Grady, F., Guha, S., Upadhyay, K., Ben-Avraham, D., Mukherjee, S., et al. Sequencing an Ashkenazi reference panel supports population-targeted personal genomics and illuminates Jewish and European origins. Nature communications 5, 1–9 (2014).

35. Porubsky, D., Ebert, P., Audano, P. A., Vollger, M. R., Harvey, W. T., Marijon, P., Ebler, J., Munson, K. M., Sorensen, M., Sulovari, A., et al. Fully phased human genome assembly without parental data using single-cell strand sequencing and long reads. Nature biotechnology 39, 302–308 (2021).

36. Byrska-Bishop, M., Evani, U. S., Zhao, X., Basile, A. O., Abel, H. J., Regier, A. A., Corvelo, A., Clarke, W. E., Musunuri, R., Nagulapalli, K., et al. High coverage whole genome sequencing of the expanded 1000 Genomes Project cohort including 602 trios. bioRxiv (2021).

37. Szustakowski, J. D., Balasubramanian, S., Kvikstad, E., Khalid, S., Bronson, P. G., Sasson, A., Wong, E., Liu, D., Wade Davis, J., Haefliger, C., et al. Advancing human genetics research and drug discovery through exome sequencing of the UK Biobank. en. Nat. Genet. 53, 942–948 (July 2021).

38. Wang, Q., Dhindsa, R. S., Carss, K., Harper, A. R., Nag, A., Tachmazidou, I., Vitsios, D., Deevi, S. V. V., Mackay, A., Muthas, D., et al. Rare variant contribution to human disease in 281,104 UK Biobank exomes. en. Nature 597, 527–532 (Sept. 2021).

39. Backman, J. D., Li, A. H., Marcketta, A., Sun, D., Mbatchou, J., Kessler, M. D., Benner, C., Liu, D., Locke, A. E., Balasubramanian, S., et al. Exome sequencing and analysis of 454,787 UK Biobank participants. en. Nature, 1–10 (Oct. 2021).

40. Wu, M. C., Lee, S., Cai, T., Li, Y., Boehnke, M. & Lin, X. Rare-variant association testing for sequencing data with the sequence kernel association test. en. Am. J. Hum. Genet. 89, 82–93 (July 2011).

41. Landrum, M. J., Lee, J. M., Riley, G. R., Jang, W., Rubinstein, W. S., Church, D. M. & Maglott, D. R. ClinVar: public archive of relationships among sequence variation and human phenotype. en. Nucleic Acids Res. 42, D980–5 (Jan. 2014).

42. Sun, C. & Medvedev, P. VarMatch: robust matching of small variant datasets using flexible scoring schemes. Bioinformatics 33, 1301–1308 (2017).

43. Hagiwara, K., Edmonson, M. N., Wheeler, D. A. & Zhang, J. indelPost: harmonizing ambiguities in simple and complex indel alignments. Bioinformatics (2021).

44. Li, H. A statistical framework for SNP calling, mutation discovery, association mapping and population genetical parameter estimation from sequencing data. Bioinformatics 27, 2987–2993 (2011).

45. Yi, R., Chang, P.-C., Baid, G. & Carroll, A. Learning from Data-Rich Problems: A Case Study on Genetic Variant Calling. arXiv preprint arXiv:1911.05151(2019).

