## Supplementary Materials for "Improving variant calling using population data and deep learning"

### Supplementary Information for: Improving variant calling using population data and deep learning

October 23, 2021

#### Supplementary Notes

##### S1 Using imputation for WGS variant calls

Imputation methods [1, 2] use population data to estimate missing genotypes and are commonly used to improve genotyping accuracy for low-coverage datasets. We applied the Minimac4 imputation method for the HG003 WGS datasets with different coverages and compared the results with DeepVariant-AF. We used the 1000Genomes GRCh38 SNV-and-indel callset as the reference panel [3].

DeepVariant-AF outperforms all imputation results when measuring the total number of SNP and indel errors. When considering variant types, DeepVariant-AF shows higher accuracy for all results but imputed DeepVariant SNP calls at 6x coverage. This suggests that the proposed population-aware variant calling framework might better utilize population information than an imputation engine for WGS datasets, even at low coverage levels.

##### S1.1 Pipeline

We first pre-processed the reference panel using Minimac3:

```
> parallel -j 30 "Minimac3 \
--refHaps ALL.chr{}.shapeit2_integrated_snvindels_v2a_27022019.GRCh38.phased.vcf.gz \
--prefix chr{} --cpus 1 --processReference" ::: $(seq 1 22)
```

We then ran imputation using Minimac4. We used files `GRCh38.rev_chrom_map` and `GRCh38.chrom_map` to convert chromosomes from format "chr1" to "1" and vice versa:

```
> DIR="dir"
> BIN="minimac4_bin"
> VCF="to_impute.vcf.gz"
> parallel -j 24 "bcftools view ${VCF} chr{} | \
bcftools annotate --rename-chrs ${DIR}/grch38_full_hla/GRCh38.rev_chrom_map \
-O z -o chr{}.vcf.gz; bcftools index chr{}.vcf.gz" ::: $(seq 1 22)
> parallel -j 24 "${BIN}/minimac4 --refHaps ${DIR}/minimac3/chr{}.m3vcf.gz \
--haps chr{}.vcf.gz --prefix chr{}.minimac4_1kgp_grch38" ::: $(seq 1 22)
> for i in $(seq 1 22); do \
ls chr${i}.minimac4_1kgp_grch38.dose.vcf.gz >> minimac4.output; done
> bcftools concat -f minimac4.output | \
bcftools annotate --rename-chrs ${DIR}/grch38_full_hla/GRCh38.chrom_map | \
bcftools view -e 'GT="0|0"' -O z -o HG003.minimac4.vcf.gz
> tabix HG003.minimac4.vcf.gz
```

#### S2 Performance on zero-frequency variants

A potential concern for population-aware variant calling models is increasing false negative rate for novel alleles. Since it is not trivial to define a set of truly novel variants in the 1000 Genomes Project, we extracted variants with zero allele frequency to investigate the impact when population information is included in a variant calling model. Using the GIAB v4.2.1 truth set, there are 32,256 (1.0%) SNPs and 3,193 (0.6%) indels that have zero allele frequency for sample HG003. We then use the zero-frequency variant set to evaluate recall of actual variant calls using hap.py [4].

We observed that the recall on zero-frequency variants underperforms the rest using all DeepVariant models, regardless of variant types and whether to utilize population information (Figure S1). With 35x reads, the recall of the population-agnostic DeepVariant model (DeepVariant) is 0.8938 for SNPs and 0.7337 for indels. The recall further decreases to 0.8813 for SNPs and 0.7142 for indels when using DeepVariant-AF. When using 21x reads, the drop in accuracy gets larger for both types of variants. This is consistent with our analysis that the population-aware DeepVariant model requires stronger evidence (higher-quality pileup images) to call zero-frequency variants, thus reducing recall. Further, the population information has a stronger influence in variant calling for low-coverage datasets. Despite the disadvantages, the negative impact on zero-frequency variants is small compared to overall error reduction.

To better understand the zero-frequency variants, we called variants using the DeepVariant PacBio model with the PrecisionFDA v2 35x HG003 reads set sequenced with the PacBio HiFi technology [5]. The recall for the zero-frequency variants improves to 0.9519 for SNPs and 0.9132 for indels. The large difference in recall/FNR indicates that many of the zero-frequency variants are hard to genotype using Illumina reads, and may not be novel mutations relative to samples in reference panels. In the future, reference panels utilizing high-quality long reads will likely provide better allele frequency estimates and improve the population-aware model performance [6–8].

### Supplementary Tables

Table S1: Variant calling and imputation results for WGS HG003. We used Minimac4 and the 1000Genomes reference panel to perform imputation. We were unable to perform imputation on the results called from Strelka2

| Coverage | Type | Method | Imputation | Recall | Precision | F1 score |
|---|---|---|---|---|---|---|
| 35x | INDEL | DeepVariant-AF | No | 0.994056 | 0.997466 | 0.995758 |
| | | DeepVariant | No | 0.993992 | 0.997365 | 0.995676 |
| | | | Yes | 0.574252 | 0.824548 | 0.677006 |
| | | GATK | No | 0.991594 | 0.991121 | 0.991357 |
| | | | Yes | 0.574036 | 0.824247 | 0.676755 |
| | | Octopus | No | 0.989744 | 0.996324 | 0.993023 |
| | | | Yes | 0.589009 | 0.830662 | 0.689269 |
| | | Strelka2 | No | 0.987627 | 0.995968 | 0.99178 |
| | SNP | DeepVariant-AF | No | 0.993929 | 0.998499 | 0.996208 |
| | | DeepVariant | No | 0.993824 | 0.998177 | 0.995996 |
| | | | Yes | 0.87932 | 0.907848 | 0.893356 |
| | | GATK | No | 0.992233 | 0.9884 | 0.990313 |
| | | | Yes | 0.879116 | 0.907008 | 0.892844 |
| | | Octopus | No | 0.988984 | 0.995357 | 0.99216 |
| | | | Yes | 0.906451 | 0.919746 | 0.91305 |
| | | Strelka2 | No | 0.992187 | 0.981123 | 0.986624 |
| 21x | INDEL | DeepVariant-AF | No | 0.984279 | 0.992219 | 0.988233 |
| | | DeepVariant | No | 0.982706 | 0.991749 | 0.987206 |
| | | | Yes | 0.573766 | 0.824062 | 0.676505 |
| | | GATK | No | 0.975544 | 0.982909 | 0.979213 |
| | | | Yes | 0.573175 | 0.822991 | 0.675734 |
| | | Octopus | No | 0.97672 | 0.99065 | 0.983636 |
| | | | Yes | 0.587183 | 0.826976 | 0.686749 |
| | | Strelka2 | No | 0.958601 | 0.989087 | 0.973605 |
| | | DeepVariant-AF | No | 0.991864 | 0.996686 | 0.994269 |
| | | DeepVariant | No | 0.991698 | 0.996023 | 0.993856 |
| | | | Yes | 0.8791 | 0.907427 | 0.893039 |
| | | GATK | No | 0.98848 | 0.987216 | 0.987847 |
| | | | Yes | 0.878593 | 0.906298 | 0.892231 |
| | | Octopus | No | 0.984764 | 0.993798 | 0.989261 |
| | | | Yes | 0.903532 | 0.916044 | 0.909745 |
| | | Strelka2 | No | 0.981011 | 0.992999 | 0.986968 |

*Continued on next page*

Table S1 – *Continued from previous page*

| **Coverage** | **Type** | **Method** | **Imputation** | **Recall** | **Precision** | **F1 score** |
|---|---|---|---|---|---|---|
| 10x | INDEL | DeepVariant-AF | No | 0.889863 | 0.958186 | 0.922762 |
| | | DeepVariant | No | 0.880882 | 0.953219 | 0.915624 |
| | | | Yes | 0.56818 | 0.817368 | 0.670366 |
| | | GATK | No | 0.865516 | 0.944581 | 0.903322 |
| | | | Yes | 0.56735 | 0.817218 | 0.669737 |
| | | Octopus | No | 0.889788 | 0.9577 | 0.922496 |
| | | | Yes | 0.574967 | 0.810379 | 0.672671 |
| | SNP | Strelka2 | No | 0.750313 | 0.96193 | 0.843045 |
| | | DeepVariant-AF | No | 0.952551 | 0.985066 | 0.968536 |
| | | DeepVariant | No | 0.950283 | 0.978971 | 0.964414 |
| | | | Yes | 0.874391 | 0.902605 | 0.888274 |
| | | GATK | No | 0.939326 | 0.976279 | 0.957446 |
| | | | Yes | 0.872958 | 0.902162 | 0.88732 |
| | | Octopus | No | 0.938012 | 0.984203 | 0.960552 |
| | | | Yes | 0.886501 | 0.898896 | 0.892655 |
| | | Strelka2 | No | 0.916328 | 0.976271 | 0.94535 |
| 6x | INDEL | DeepVariant-AF | No | 0.72068 | 0.914249 | 0.806005 |
| | | DeepVariant | No | 0.710767 | 0.902276 | 0.795153 |
| | | | Yes | 0.555917 | 0.804138 | 0.657376 |
| | | GATK | No | 0.691391 | 0.898816 | 0.781576 |
| | | | Yes | 0.554787 | 0.805245 | 0.656954 |
| | | Octopus | No | 0.748593 | 0.905851 | 0.819748 |
| | | | Yes | 0.544975 | 0.770679 | 0.638467 |
| | | Strelka2 | No | 0.500348 | 0.935336 | 0.651945 |
| | SNP | DeepVariant-AF | No | 0.82965 | 0.962828 | 0.891292 |
| | | DeepVariant | No | 0.82624 | 0.948907 | 0.883335 |
| | | | Yes | 0.859202 | 0.891795 | 0.875195 |
| | | GATK | No | 0.80301 | 0.951194 | 0.870843 |
| | | | Yes | 0.855737 | 0.891706 | 0.873351 |
| | | Octopus | No | 0.834285 | 0.953795 | 0.890047 |
| | | | Yes | 0.842877 | 0.856744 | 0.849754 |
| | | Strelka2 | No | 0.753403 | 0.917202 | 0.827272 |

Table S2: Variant calling results for WES HG003 (Oslo dataset)

| **Type** | **Caller** | **False negatives** | **False positives** | **Recall** | **Precision** | **F1 score** |
|---|---|---|---|---|---|---|
| INDEL | DeepVariant-AF | 56 | 30 | 0.965389 | 0.981343 | 0.973301 |
| | DeepVariant | 61 | 35 | 0.962299 | 0.978234 | 0.970201 |
| | GATK | 113 | 182 | 0.930161 | 0.893255 | 0.911334 |
| | Octopus | 82 | 71 | 0.94932 | 0.956996 | 0.953143 |
| | Strelka2 | 104 | 80 | 0.935723 | 0.950403 | 0.943006 |
| SNP | DeepVariant-AF | 270 | 100 | 0.990198 | 0.996347 | 0.993263 |
| | DeepVariant | 279 | 112 | 0.989871 | 0.995909 | 0.992881 |
| | GATK | 283 | 365 | 0.989726 | 0.986784 | 0.988252 |
| | Octopus | 456 | 213 | 0.983445 | 0.99219 | 0.987798 |
| | Strelka2 | 379 | 138 | 0.98624 | 0.994945 | 0.990574 |

Table S3: Variant calling results for WES HG003 (IDT dataset)

| **Type** | **Caller** | **False negatives** | **False positives** | **Recall** | **Precision** | **F1 score** |
|---|---|---|---|---|---|---|
| INDEL | DeepVariant-AF | 28 | 14 | 0.975779 | 0.987952 | 0.981827 |
| | DeepVariant | 31 | 21 | 0.973183 | 0.981974 | 0.977559 |
| | GATK | 74 | 315 | 0.935986 | 0.780181 | 0.851011 |
| | Octopus | 52 | 81 | 0.955017 | 0.933168 | 0.943966 |
| | Strelka2 | 63 | 143 | 0.945502 | 0.886328 | 0.914959 |
| SNP | DeepVariant-AF | 258 | 131 | 0.989845 | 0.994818 | 0.992326 |
| | DeepVariant | 282 | 135 | 0.988901 | 0.994656 | 0.99177 |
| | GATK | 266 | 433 | 0.98953 | 0.98306 | 0.986285 |
| | Octopus | 437 | 216 | 0.9828 | 0.991418 | 0.98709 |
| | Strelka2 | 365 | 141 | 0.985634 | 0.994401 | 0.98999 |

Table S4: Variant calling accuracy on common (allele frequency $> 0.01$) variants. Dataset: HG003 35x WGS

| **Type** | **Caller** | **False negatives** | **False positives** | **Recall** | **Precision** | **F1 score** |
|---|---|---|---|---|---|---|
| INDEL | DeepVariant-AF | 2055 | 1361 | 0.995829 | 0.997351 | 0.9965894 |
| | DeepVariant | 2179 | 1345 | 0.995577 | 0.997382 | 0.9964787 |
| | GATK | 3465 | 2565 | 0.992967 | 0.994999 | 0.9939820 |
| | Strelka2 | 5344 | 2104 | 0.989153 | 0.995855 | 0.9924927 |
| SNP | DeepVariant-AF | 10209 | 4401 | 0.996835 | 0.998634 | 0.9977337 |
| | DeepVariant | 10979 | 4415 | 0.996597 | 0.998629 | 0.9976120 |
| | GATK | 16202 | 13630 | 0.994978 | 0.995773 | 0.9953753 |
| | Strelka2 | 29972 | 3925 | 0.990709 | 0.998774 | 0.9947252 |

Table S5: Variant calling accuracy on rare (allele frequency $\leq 0.01$) variants. Dataset: HG003 35x WGS

| **Type** | **Caller** | **False negatives** | **False positives** | **Recall** | **Precision** | **F1 score** |
|---|---|---|---|---|---|---|
| INDEL | DeepVariant-AF | 949 | 207 | 0.919583 | 0.983799 | 0.9506077 |
| | DeepVariant | 866 | 306 | 0.926616 | 0.976500 | 0.9509042 |
| | GATK | 635 | 2221 | 0.946191 | 0.859350 | 0.9006821 |
| | Strelka2 | 998 | 1059 | 0.915431 | 0.925794 | 0.9205833 |
| SNP | DeepVariant-AF | 4033 | 678 | 0.943084 | 0.992656 | 0.9672353 |
| | DeepVariant | 3876 | 1750 | 0.945300 | 0.981345 | 0.9629853 |
| | GATK | 3980 | 25226 | 0.943832 | 0.784869 | 0.8570418 |
| | Strelka2 | 4338 | 3079 | 0.938780 | 0.967704 | 0.9530226 |

Table S6: Software used in the experiments

| | |
|---|---|
| DeepVariant [9] | 1.1 |
| GATK [10] | 4.2.0.0 |
| Octopus [11] | 0.7.2 |
| Strelka2 [12] | 2.9.2 |
| Minimac3 [1] | 2.0.1 |
| Minimac4 [1] | 1.0.2 |
| GNU Parallel [13] | 20200322 |

### Supplementary Figures

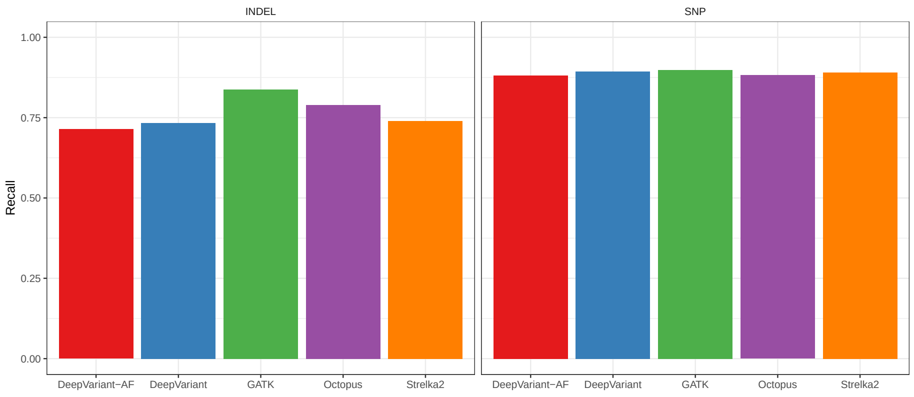


Figure S1: Recall for HG003 GIAB v4.2.1 variants that have zero allele frequency in the 1000 Genomes Project

### References

1. Das, S., Forer, L., Schönherr, S., Sidore, C., Locke, A. E., Kwong, A., Vrieze, S. I., Chew, E. Y., Levy, S., McGue, M., *et al.* Next-generation genotype imputation service and methods. *Nature genetics* **48,** 1284–1287 (2016).

2. Browning, B. L., Zhou, Y. & Browning, S. R. A one-penny imputed genome from next-generation reference panels. *The American Journal of Human Genetics* **103,** 338–348 (2018).

3. Lowy-Gallego, E., Fairley, S., Zheng-Bradley, X., Ruffier, M., Clarke, L., Flicek, P., Consortium, 1. G. P., *et al.* Variant calling on the GRCh38 assembly with the data from phase three of the 1000 Genomes Project. *Wellcome Open Research* **4** (2019).

4. Krusche, P., Trigg, L., Boutros, P. C., Mason, C. E., Francisco, M., Moore, B. L., Gonzalez-Porta, M., Eberle, M. A., Tezak, Z., Lababidi, S., *et al.* Best practices for benchmarking germline small-variant calls in human genomes. *Nature biotechnology* **37,** 555–560 (2019).

5. Olson, N. D., Wagner, J., McDaniel, J., Stephens, S. H., Westreich, S. T., Prasanna, A. G., Johanson, E., Boja, E., Maier, E. J., Serang, O., *et al.* precisionFDA Truth Challenge V2: Calling variants from short-and long-reads in difficult-to-map regions. *bioRxiv* (2020).

6. Ebert, P., Audano, P. A., Zhu, Q., Rodriguez-Martin, B., Porubsky, D., Bonder, M. J., Sulovari, A., Ebler, J., Zhou, W., Mari, R. S., *et al.* Haplotype-resolved diverse human genomes and integrated analysis of structural variation. *Science* **372** (2021).

7. De Coster, W., Weissensteiner, M. H. & Sedlazeck, F. J. Towards population-scale long-read sequencing. *Nature Reviews Genetics,* 1–16 (2021).

8. Beyter, D., Ingimundardottir, H., Oddsson, A., Eggertsson, H. P., Bjornsson, E., Jonsson, H., Atlason, B. A., Kristmundsdottir, S., Mehringer, S., Hardarson, M. T., *et al.* Long-read sequencing of 3,622 Icelanders provides insight into the role of structural variants in human diseases and other traits. *Nature Genetics* **53,** 779–786 (2021).

9. Poplin, R., Chang, P.-C., Alexander, D., Schwartz, S., Colthurst, T., Ku, A., Newburger, D., Dijamco, J., Nguyen, N., Afshar, P. T., *et al.* A universal SNP and small-indel variant caller using deep neural networks. *Nature biotechnology* **36,** 983–987 (2018).

10. Van der Auwera, G. A. & O'Connor, B. D. *Genomics in the Cloud: Using Docker, GATK, and WDL in Terra* (O'Reilly Media, 2020).

11. Cooke, D. P., Wedge, D. C. & Lunter, G. A unified haplotype-based method for accurate and comprehensive variant calling. *Nature biotechnology,* 1–8 (2021).

12. Kim, S., Scheffler, K., Halpern, A. L., Bekritsky, M. A., Noh, E., Källberg, M., Chen, X., Kim, Y., Beyter, D., Krusche, P., *et al.* Strelka2: fast and accurate calling of germline and somatic variants. *Nature methods* **15,** 591–594 (2018).

13. Tange, O. *GNU Parallel 2018* ISBN: 9781387509881. `https://doi.org/10.5281/zenodo.1146014` (Ole Tange, Mar. 2018).
